# Surface Hydrophilicity Promotes Bacterial Twitching Motility

**DOI:** 10.1101/2024.03.23.586342

**Authors:** Megan T. O’Hara, Tori M. Shimozono, Keane J. Dye, David Harris, Zhaomin Yang

**Affiliations:** Department of Biological Sciences, Virginia Tech, Blacksburg, Virginia, USA; Uniformed Services University of the Health Sciences, Bethesda, Maryland, USA; Department of Biochemistry, Virginia Tech, Blacksburg, Virginia, USA

**Keywords:** Twitching motility, *Pseudomonas aeruginosa*, *Acinetobacter*, bile salts, detergents, surface property, hydrophilicity

## Abstract

Twitching motility is a form of bacterial surface translocation powered by the type IV pilus (T4P). It is frequently analyzed by interstitial colony expansion between agar and the polystyrene surfaces of Petri dishes. In such assays, the twitching motility of *Acinetobacter nosocomialis* was observed with MacConkey but not Luria-Bertani (LB) agar media. One difference between these two media is the presence of bile salts as a selective agent in MacConkey but not in LB. Here, we demonstrate that the addition of bile salts to LB allowed *A. nosocomialis* to display twitching. Similarly, bile salts enhanced the twitching of *Acinetobacter baumannii* and *Pseudomonas aeruginosa* in LB. These observations suggest that there is a common mechanism whereby bile salts enhance bacterial twitching and promote interstitial colony expansion. Bile salts disrupt lipid membranes and apply envelope stress as detergents. Surprisingly, their stimulatory effect on twitching appears not to be related to a bacterial physiological response to stressors. Rather it is due to their ability to alter the physicochemical properties of a twitching surface. We observed that while other detergents promoted twitching like bile salts, stresses applied by antibiotics, including the outer membrane-targeting polymyxin B, did not enhanced twitching motility. More importantly, bacteria displayed increased twitching on hydrophilic surfaces such as those of glass and tissue culture-treated polystyrene plastics, and bile salts no longer stimulated twitching on these surfaces. Together, our results show that altering the hydrophilicity of a twitching surface significantly impacts T4P functionality.

**Importance:** The bacterial type IV pilus (T4P) is a critical virulence factor for many medically important pathogens, some of which are prioritized by the World Health Organization for their high levels of antibiotic resistance. The T4P is known to propel bacterial twitching motility, providing a convenient assay for T4P functionality. Here, we show that bile salts and other detergents augment the twitching of multiple bacterial pathogens. We identified the underlying mechanism as the alteration of surface hydrophilicity by detergents. Consequently, hydrophilic surfaces such as those of glass or plasma-treated polystyrene promote bacterial twitching, bypassing the requirement for detergents. The implication is that surface properties, such as those of tissues and medical implants, significantly impact the functionality of bacterial T4P as a virulence determinant. This offers valuable insights for developing countermeasures against the colonization and infection by bacterial pathogens of critical importance to human health on a global scale.

## Introduction

Twitching motility is a form of non-flagellated bacterial locomotion that allows bacteria to move on or between solid surfaces (1–4). It is powered by the bacterial type IV pilus (T4P) which can be assembled and disassembled by the supramolecular T4P machinery (T4PM) (1, 5–7). The current model proposes that it is the recurrent cycles of T4P assembly and disassembly, or extension and retraction, that powers this form of bacterial surface motility (8, 9). The T4PM assembles the long T4P filament that protrudes from a cell into its surroundings. When the tip of an extended T4P attaches to a solid substratum, the retraction of the T4P by the T4PM moves a bacterium toward the point of attachment. This translocation of bacterial cells on or between solid surfaces, results in bacterial twitching motility.

Of relevance to human health, the T4P plays a crucial role in the pathogenesis of many important bacterial pathogens (10–15). These include *Pseudomonas aeruginosa* and *Acinetobacter baumannii,* both on the list of priority pathogens per the World Health Organization (WHO) (16). One of the primary functions of the T4P as a virulence factor is for adherence to human cells or tissues to initiate colonization and invasion (4, 17, 18).

*Acinetobacter nosocomialis,* a close relative of *A. baumannii,* is an opportunistic pathogen primarily causing nosocomial or hospital-acquired infections (19). The *A. nosocomialis* M2 strain has been used as a model for studies of *Acinetobacter* pathogenesis and T4P functionality (18, 20–22). Despite the lack of flagella and the acineto-or non-motile designation for this genus, many *Acinetobacter* species are, in fact, motile by T4P-dependent twitching motility (23–26). As such, twitching motility provides a convenient assay for investigating the functionality of the bacterial T4P in these medically important pathogens.

Twitching motility is routinely analyzed by observing interstitial colony expansion between the lower surface of solidified nutrient agar and that of plastic Petri dishes made of polystyrene (25, 27). Such stab assays involve the inoculation of the interstitial space by stabbing through the agar, and this method has been used for the identification of T4P or *pil* genes by the isolation of *P. aeruginosa* mutants that were defective in twitching motility (3). T4P genes encode the core components of the T4PM and their functions in twitching motility are conserved among *P. aeruginosa* and many Gram-negative and Gram-positive bacteria (2, 10, 28–31). These include PilA, the major pilin, as well as PilB, the T4P extension ATPase, and PilT, the T4P retraction ATPase. Along with other T4P proteins, the PilB and the PilT ATPases polymerize and depolymerize pilins into or from the T4P filament, respectively. Bacterial translocation by twitching motility over distances longer than the length of an extended pilus depends on the dynamic nature of T4P assembly and disassembly coordinated by the T4PM (8, 9).

The regulation of bacterial motility by environmental cues has been studied most extensively in flagellated bacteria (32–35). Besides chemotactic responses (36), the biogenesis of bacterial flagella is modulated through gene expression and flagellar assembly by signals such as nutrient and surface availability (32–35). In addition, an alternative sigma factor, which is responsive to envelope and other environmental stressors, transcriptionally regulates the expression of flagellar genes in many bacteria (37–39). Although the T4P has been investigated to a lesser extent, there is clear evidence that its biogenesis and function are influenced by regulatory mechanisms and environmental factors. In many T4P or *pil* gene clusters, there are conserved two-component systems including PilS and PilR (2, 31). In selected organisms, these regulators have been demonstrated to affect the expression of T4P genes (40–43). Signals of both chemical and physical nature are known to influence T4P-mediated motility (44–46). For example, lactate can induce PilT-dependent T4P retraction in *Neisseria meningitidis* whereas both temperature and blue light were shown to influence bacterial twitching motility (47, 48). It was observed previously that *A. nosocomialis* exhibited twitching motility only with MacConkey but not with Luria-Bertani (LB) agar media on polystyrene Petri dishes (22, 49). In our current study, we investigated the underlying reasons for the observed differences in bacterial twitching between these two media. We determined that bile salts are the key component that allows *A. nosocomialis* to twitch in MacConkey media. This is because the addition of bile salts to LB allowed *A. nosocomialis* to twitch to a similar extent as with MacConkey. We also observed similar stimulatory effects of bile salts on the twitching of *P. aeruginosa* and *A. baumannii*. Bile salts are anionic detergents that can apply membrane stress to bacteria (50–53). Our results further demonstrate that other detergents likewise can enhance bacterial twitching on polystyrene surfaces. Antibiotics, including the outer membrane-targeting polymyxin B, do not increase *P. aeruginosa* twitching motility. This suggests that the mechanism for the stimulatory effect of bile salts is unlikely related to a bacterial response to the presence of a general or envelope stressor. Instead, we suspected that bile salts and other detergents increased the hydrophilicity of polystyrene surfaces, and it was this increase in hydrophilicity that promoted bacterial twitching. Indeed, we observed that glass surfaces, which are more hydrophilic than polystyrene, significantly promoted twitching motility. In contrast, increasing the hydrophobicity of glass surfaces by coating attenuated twitching. Like glass, plasma-treatment of polystyrene surfaces is known to increase their hydrophilicity for culturing tissues or cells. We observed that tissue culture (TC)-treated polystyrene surfaces significantly increased bacterial twitching. Moreover, the addition of bile salts no longer stimulated twitching on glass or TC-treated polystyrene surfaces. Our results here suggest that bacterial pathogens may have evolved mechanisms to differentially interact with surfaces that have varying physicochemical properties to optimize host recognition, colonization, and infections.

## Results

### Bile salts enable *Acinetobacter* twitching motility in stab assays

It has been reported in the literature that *A. nosocomialis* displays twitching motility with MacConkey, but not LB agar, in stab assays (22, 49). In these assays, bacterial cells are stab-inoculated through the agar to form an interstitial colony between the Petri dish and the agar media (25, 27). After a period of incubation, the size of an interstitial colony can be measured to quantify twitching motility. As shown in Figure 1A, the *A. nosocomialis* M2 strain shows clear twitching with MacConkey, but not with LB agar. These results confirmed that MacConkey but not LB media allows *A. nosocomialis* to display twitching motility as observed previously (22, 49).

**Figure 1.**
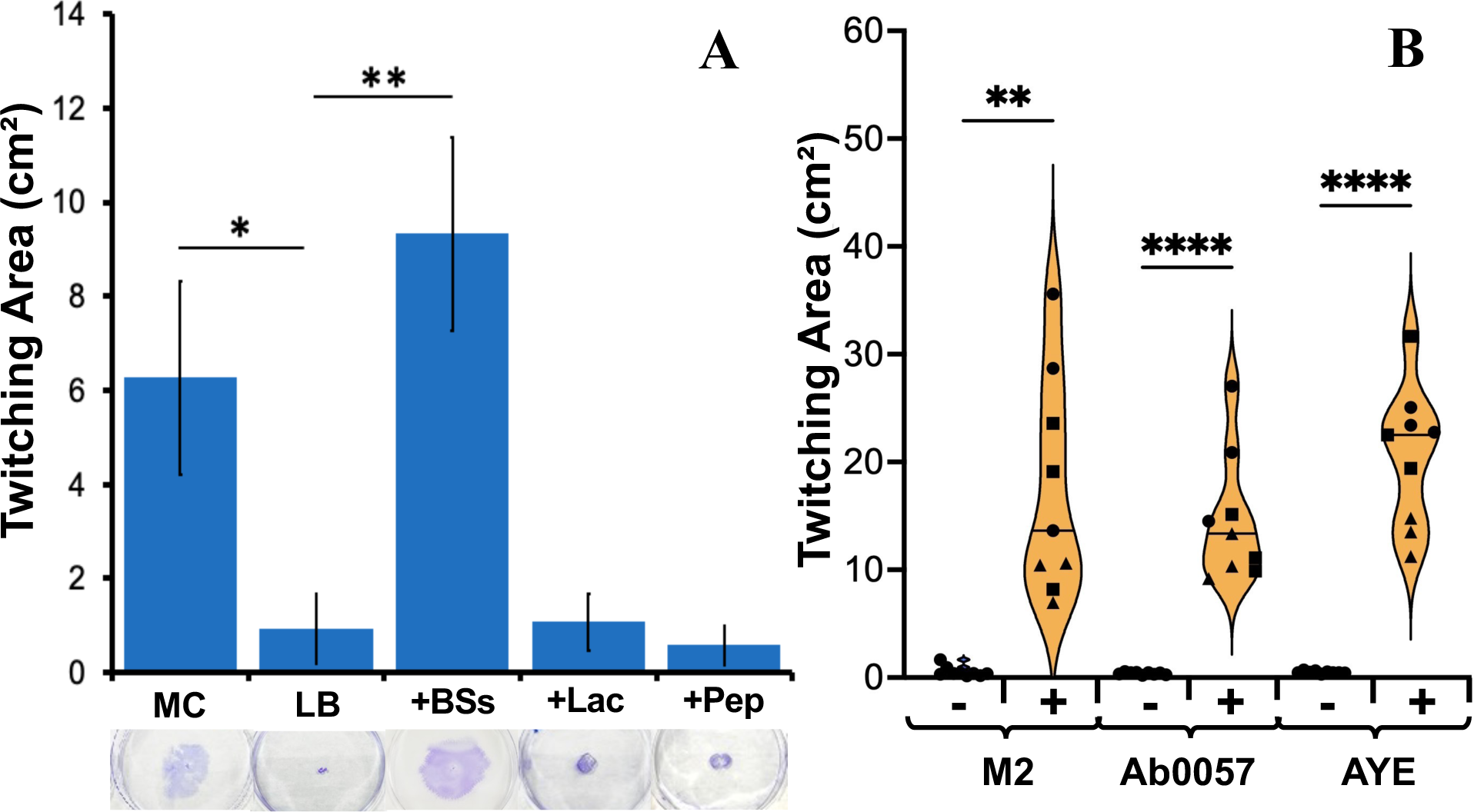
Bile salts enable *Acinetobacter* to twitch. **A.** Bile salts allow *A. nosocomialis* M2 to twitch in LB media. The twitching motility of *A. nosocomialis* M2 was analyzed with MacConkey (MC) or Luria-Bertani (LB) media without or with 0.5% bile salts (+BSs), 1% lactose (+Lac), or 2% peptone (+Pep) with standard polystyrene Petri dishes as described in Materials and Methods. Data shown are the averages from three biological experiments each performed in triplicate. **B.** Bile salts provoke *A. baumannii* twitching in LB media. *A. baumannii* strains AB0057 and AYE were analyzed for twitching motility in LB agar without (-) or with (+) 0.5% bile salts on standard polystyrene Petri dishes with *A. nosocomialis* (M2) as a control. Data shown are from three biological experiments, represented by different symbols, each performed in triplicate. Single (*), double (**), and quadruple (****) asterisks indicate two values are statistically different with P<0.05, P<0.01, and P<0.0001 by the Student’s T-test, respectively.

We compared the composition of these two commonly used bacterial growth media (Table S1). Notwithstanding their commonalities, LB lacks peptone, lactose, and bile salts that are present in MacConkey. Peptone is a proteinous nutrient source and lactose is a carbon and energy source. Bile salts are cholesterol derivatives with aliphatic side chains (51, 54) that regulate various biological processes in vertebrates and their microbiomes (50, 54–57). The amphipathic nature of bile salts allows them to interact with and disrupt membranes, resulting in envelope stress in bacteria as detergents (51, 52). Both Gram-positive and Gram-negative bacteria can respond to the presence of bile salts, leading to changes in gene expression and cellular physiology (52, 58, 59). Generally, bile salts are more inhibitory of Gram-positive bacteria because of the lack of an outer membrane. Thus, they are included in MacConkey as a selective agent against Gram-positive bacteria in favor of enteric bacteria.

We supplemented LB agar with peptone, lactose, or bile salts at the same concentration present in MacConkey agar to determine if one of these could enable *A. nosocomialis* to twitch in LB. As shown in Figure 1A, the addition of neither lactose nor peptone led to any discernible twitching motility in *A. nosocomialis*. In contrast, the supplementation of bile salts resulted in *A. nosocomialis* twitching motility in LB comparable to what was observed with MacConkey agar. These results indicated that bile salts are the component that specifically stimulates twitching motility of *A. nosocomialis* as analyzed by stab assays with polystyrene Petri dishes.

Twitching motility has been observed in *A. baumannii* (24, 60), a closely related *Acinetobacter* species and a WHO priority pathogen (16). The twitching motility of this bacterium is similarly noted in MacConkey agar as analyzed by a similar assay (22, 49). However, variable motility phenotypes were observed with LB media for different clinical isolates (60), suggesting an effect of media composition on *A. baumannii* twitching. We tested two *A. baumannii* strains, AYE and AB0057 (61, 62), in LB supplemented with bile salts in comparison with *A. nosocomialis* M2. As shown in Figure 1B, while none of these strains displayed twitching motility with the LB agar, supplementation of bile salts elicited twitching motility of both *A. baumannii* strains similarly to *A. nosocomialis*. These results suggest that the stimulatory effects of bile salts on twitching motility is a more general phenomenon in the *Acinetobacter* genus.

### Bile salts stimulated *P. aeruginosa* twitching motility

The above observations prompted us to investigate if bile salts enhanced bacterial twitching in other bacteria. *P. aeruginosa*, another WHO priority pathogen (16), has been used as a model for studies of bacterial twitching (8, 63, 64). Its twitching motility has been routinely analyzed using stab assays with LB instead of MacConkey agar plates (65–68). *P. aeruginosa* PAO1, a frequently used laboratory strain, exhibits twitching motility in LB agar (69, 70). However, the addition of bile salts to LB significantly increased its twitching motility (Figure 2A). Further, we examined the dose response of PAO1 twitching to bile salts. As shown in Figure 2B, the stimulation of twitching motility shows concentration dependency, with a plateau between 0.1% and 0.4% of bile salts. At higher concentrations, bile salts start to inhibit *P. aeruginosa* growth and reduce its twitching motility in this assay (data not shown). We additionally examined the twitching motility of PA14, another commonly used *P. aeruginosa* strain in the literature (71–73). It was observed that the twitching motility of PA14 was stimulated by bile salts in LB media like that of PAO1 (Figure S1). These results demonstrate that the stimulatory effect of bile salts on twitching is applicable to both *Acinetobacter* species and *P. aeruginosa* isolates. For the remainder of this study, we mostly used *P. aeruginosa* PAO1 as the model organism to investigate the mechanisms by which bile salts stimulate bacterial twitching motility.

**Figure 2.**
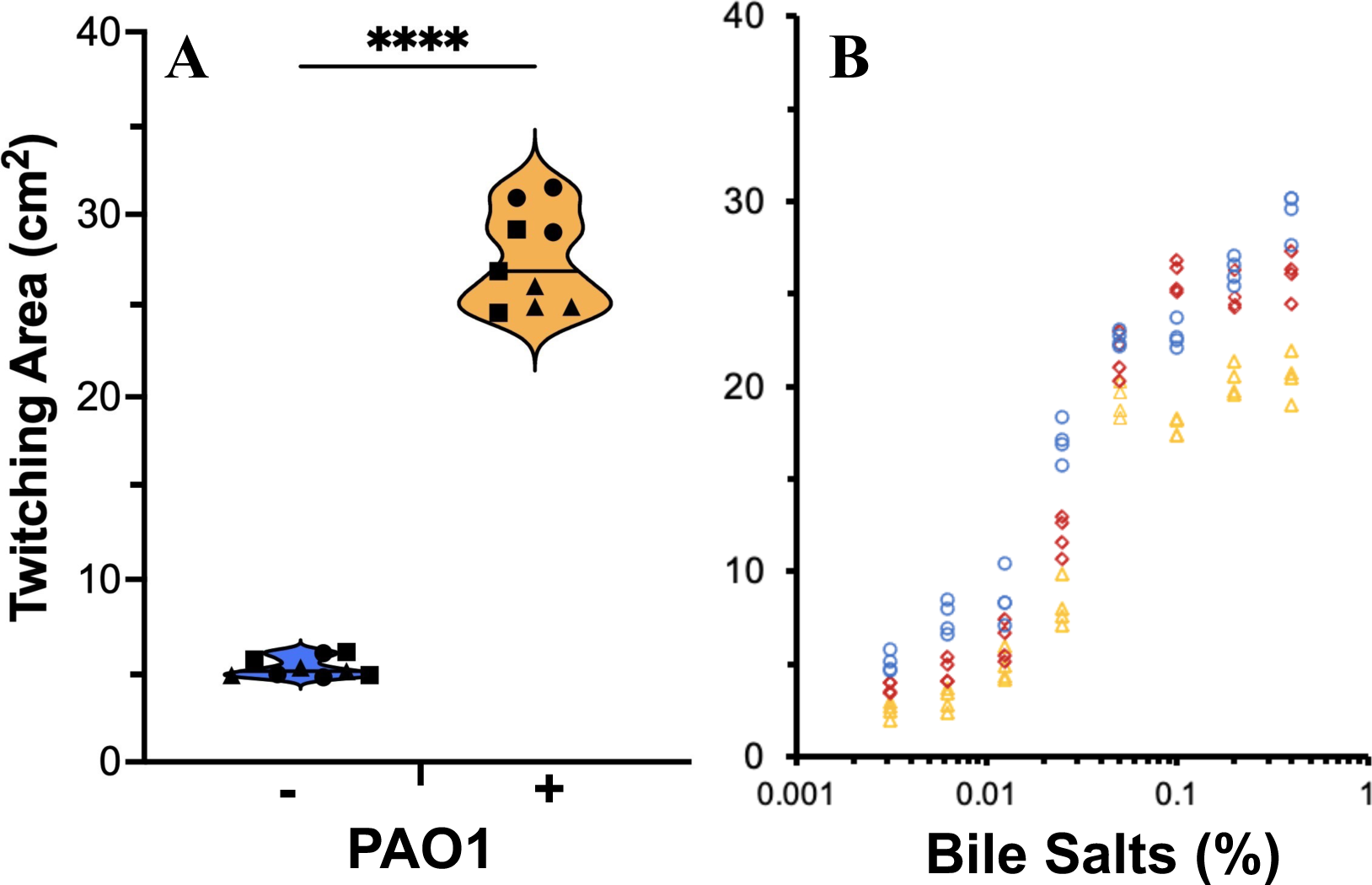
Bile salts enhance *P. aeruginosa* twitching motility. **A.** Bile salts increase *P. aeruginosa* twitching. Twitching motility of PAO1 was analyzed without (-) or with (+) 0.5% bile salts as in Figure 1B with data similarly presented. Quadruple asterisks (****) indicates values that are statistically different with P<0.0001 by the Student’s T-test. **B.** Dose effect of bile salts on PAO1 twitching. PAO1 twitching was analyzed with the standard polystyrene Petri dish protocol as in (A) with varying concentrations (w/v %) of bile salts as indicated. Data presented in both panels are from three biological experiments each performed in triplicate. Data points from the same experiment are represented by the same symbols in color and shape.

### Detergents stimulate bacterial twitching

Bile salts, produced from cholesterol metabolism, are anionic detergents (50–52, 55). They are known to apply membrane or envelope stress in bacteria (51, 52, 74). It is possible that bile salts function as a detergent to apply envelope or general stress to cells and it was the cellular stress response that underlies the stimulatory effects of bile salts on bacterial twitching motility. To examine this possibility, we investigated the effect of other detergents on the twitching motility of *P. aeruginosa.* To avoid the complications between growth inhibition and twitching motility, we determined the maximum non-inhibitory concentrations of detergents experimentally (Table S2) to guide their use in our twitching motility assays. For this experiment, we supplemented the LB media with the anionic detergent sodium dodecyl sulfate (SDS), or the non-ionic detergents Triton X-100 and Triton X-114 (Table S2). As shown in Figure 3A, all the detergents examined, whether anionic or non-ionic, significantly stimulated the twitching motility of *P. aeruginosa* much like bile salts. These results support the notion that the promotional effects of bile salts on twitching are related to their amphipathic properties as detergents.

**Figure 3.**
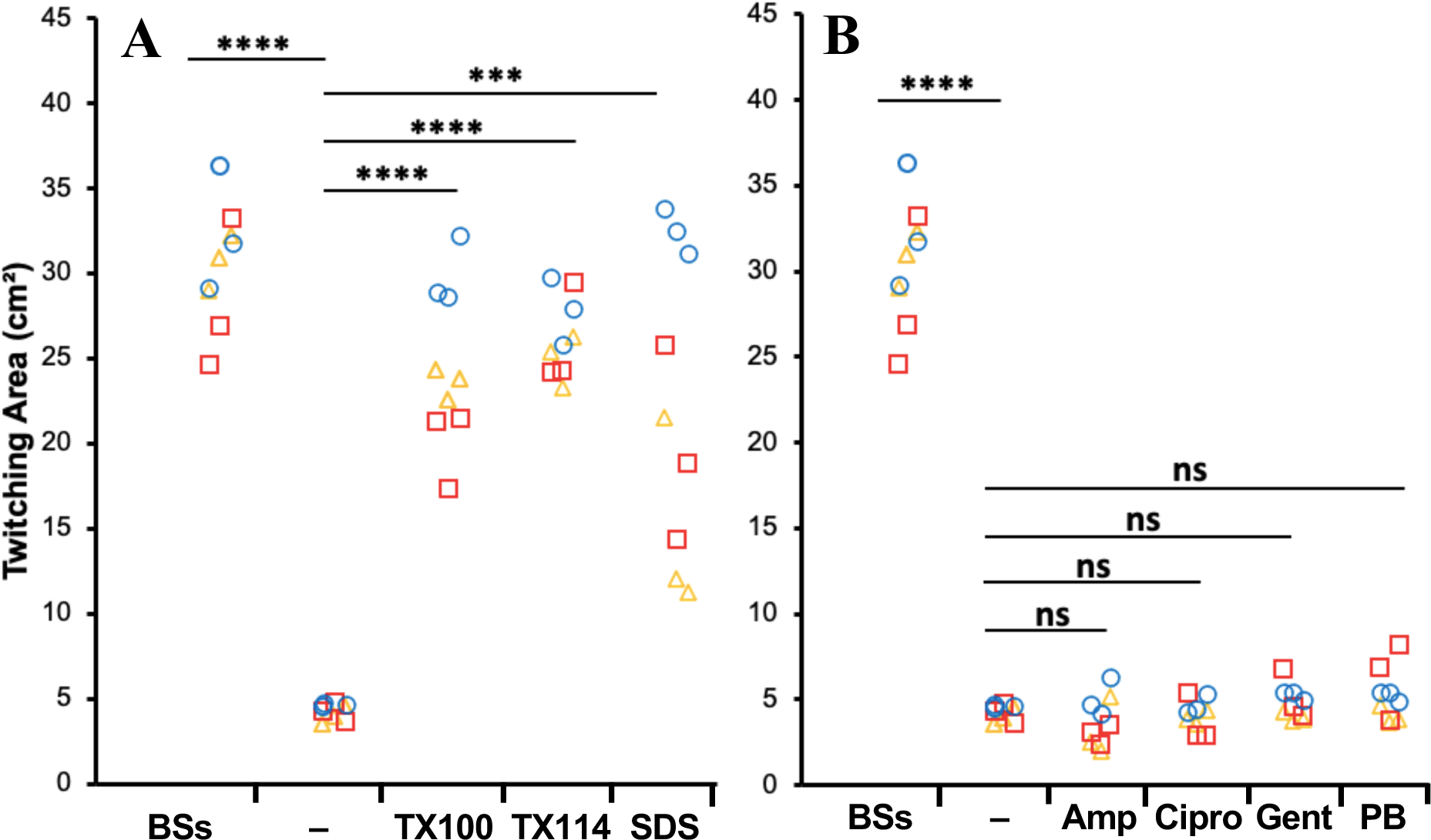
Detergents, but not antibiotics, promote *P. aeruginos*a twitching. **A.** Effects of detergents. PAO1 twitching was analyzed with the standard Petri dish protocol as in Figure 2A with LB agar without modification (-) or with bile salts (BSs) (5 mg/ml), Triton X-100 (TX100) (75 μg/ml), Triton X-114 (TX114) (75 μg/ml), or sodium dodecyl sulfate (SDS) (850 μg/ml). **B.** Effects of antibiotics. PAO1 twitching was analyzed as in (A) with ampicillin (Amp) (313 ng/ml), ciprofloxacin (Cipro) (31 ng/ml), gentamicin (Gent) (31 ng/ml), or polymyxin B (PB) (313 ng/ml). Data presented in both panels are from three biological experiments each performed in triplicate. Data points from the same experiment are represented by the same symbols in color and shape. Triple (***) and quadruple (****) asterisks indicate two values are statistically different with P<0.001 and P<0.0001 by the Student’s T-test, respectively. Antibiotics resulted in values that are not significantly (ns) different with P>0.05.

The stimulatory effects of bile salts and other detergents on twitching motility could be explained by a physiological response of a bacterium to envelope stress applied by these amphipathic molecules (51, 52, 74, 75) or a general stress response to various environmental stressors (76). To investigate this, we tested antibiotics with different modes of action at their maximum non-inhibitory concentrations as stressors. These included ampicillin, gentamicin, and ciprofloxacin, which target cell wall biosynthesis, ribosome function and DNA topology, respectively. We first determined the maximum non-inhibitory concentrations of these antibiotics by testing their effect on *P. aeruginosa* growth at different concentrations (Table S3). We then tested these antibiotics at their respective maximum non-inhibitory concentrations for their effect on *P. aeruginosa* twitching. As shown in Figure 3B, none of these above antibiotics affected *P. aeruginosa* twitching motility significantly. This suggested that the stimulation of twitching by detergents was unlikely the result of a physiological response to general stressors.

Moreover, we tested the effect of polymyxin B, which applies envelope stress, as do bile salts, by targeting the outer membrane of Gram-negative bacteria (53). Somewhat unexpectedly, this antibiotic showed no stimulatory effect on *P. aeruginosa* twitching (Figure 3B). These results suggested that the observed stimulation of twitching motility by bile salts and other detergents (Figures 2 and 3A) might not be related to a response to general or envelope stress.

### Glass enhances *P. aeruginosa* twitching in comparison to polystyrene

Detergents such as bile salts are amphipathic molecules with both polar and non-polar moieties (50, 54, 56, 57, 77). As such, they can change the physicochemical properties of a surface (50, 78–80). In stab assays for twitching motility, bacteria cells translocate in the interstitial space between the solidified agar media and the hydrophobic surface of a polystyrene Petri dish. We considered the possibility that bile salts in a growth media may interact with the hydrophobic surface of the polystyrene Petri dishes to alter its physicochemical properties. Such interactions may allow bile salts to make the polystyrene surface more hydrophilic to possibly facilitate twitching motility. In comparison with polystyrene, glass Petri dishes present a more hydrophilic surface. We therefore examined *P. aeruginosa* twitching with LB media using glass in comparison with polystyrene Petri plates. As shown in Figure S2, *P. aeruginosa* was observed to twitch significantly more on glass Petri dishes than polystyrene ones in LB without the addition of bile salts. These results are consistent with the proposition that surface hydrophobicity or hydrophilicity plays crucial roles in bacterial twitching.

### Bile salts do not enhance *P. aeruginosa* twitching on glass surfaces

We reproduced the above observation on glass (Figure S2) with a modified twitching assay where a glass or a polystyrene microscope slide was used as the twitching surface (see Materials and Methods). In this assay, the slides were cleaned and sterilized before they were placed in a polystyrene Petri dish. Molten LB agar media was then poured into the Petri dish. Twitching motility was analyzed as before, except that the incubation time was shortened to limit the twitching zone to be within the boundaries of the width of the microscope slide. As shown in Figure 4A, PAO1 twitched significantly more on glass slides than on polystyrene ones as was observed with Petri dishes (Figure S2). As expected, the addition of bile salts significantly stimulated twitching on the polystyrene slide (Figure 4A). In contrast, the supplementation of bile salts showed no promotional effect on twitching with the glass slide (Figure 4A). A *pilA* mutant, which is non-piliated, was used as the non-twitching control, and it showed no twitching motility on all surfaces with or without bile salts (Figure 4). It is also noteworthy that the twitching motility on the polystyrene slide in the presence of bile salts showed no statistical difference from that on the glass slides with or without bile salts. These results indicate that surface hydrophilicity likely enhances twitching motility, and the effects of bile salts on twitching could be attributed to their ability to change a hydrophobic surface to a more hydrophilic one. This is consistent with the observation that the stimulatory effect of bile salts is no longer observed on the more hydrophilic glass surface in contrast with polystyrene ones.

**Figure 4.**
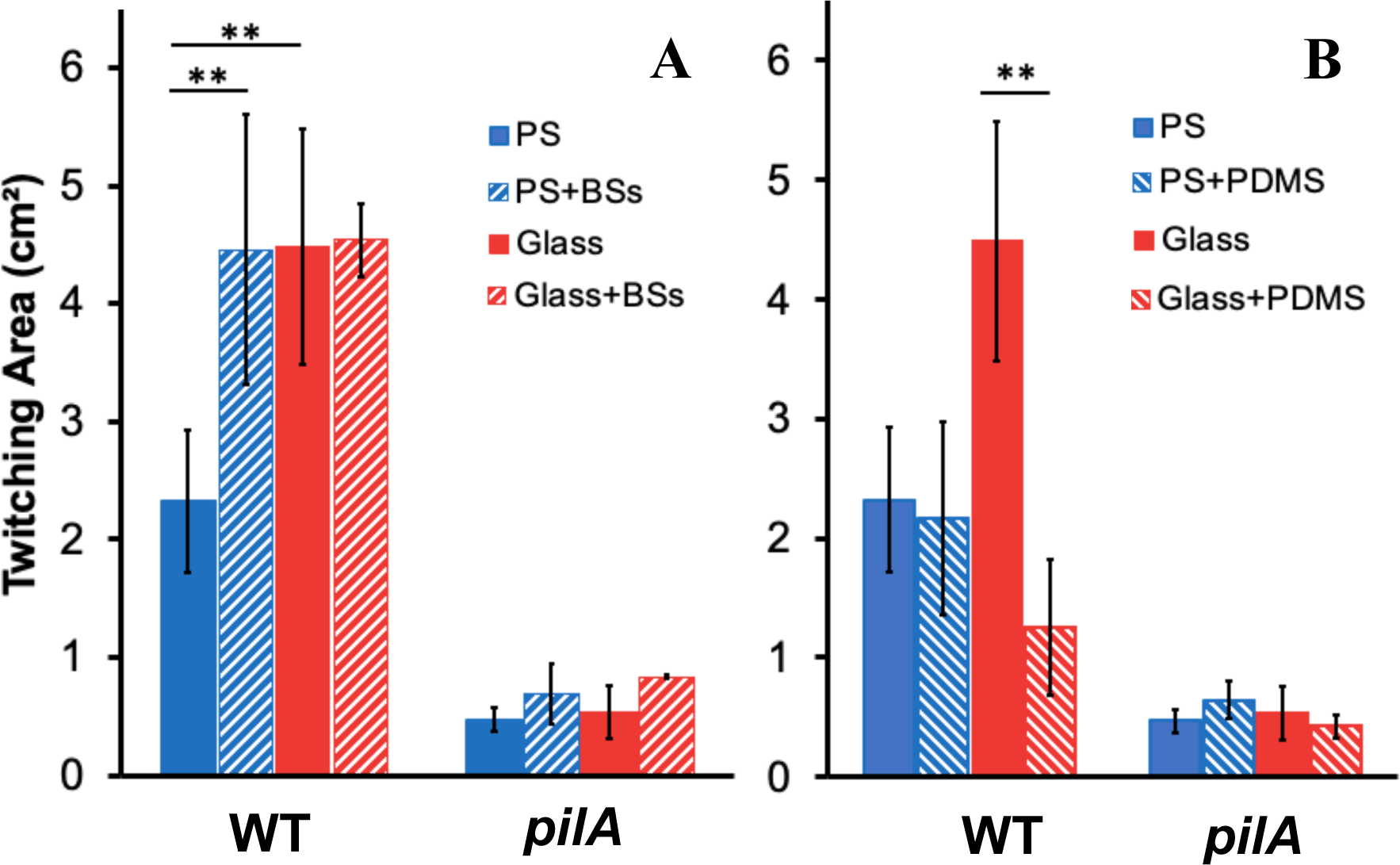
Glass surfaces increase *P. aeruginosa* twitching motility. **A.** Glass surfaces stimulate PAO1 twitching motility and eliminate the stimulatory effect of bile salts. The twitching motility of PAO1 and its isogenic *pilA* mutant were analyzed with polystyrene (PS) or glass microscope slides in LB without or with 0.5% bile salts (BSs) after 18 hours of incubation (see Materials and Methods). **B.** Hydrophobic coating of glass reduces twitching motility. Experiments were performed as in A, except that the microscope slides were coated without or with polydimethylsiloxane (PDMS) (see Materials and Methods). Data presented in both panels are from three biological experiments each performed in triplicate. Double asterisks (**) indicate two values are statistically different with P<0.01 by the Student’s T-test.

### Increase in hydrophobicity of glass surfaces reduces *P. aeruginosa* twitching

Next, we modified the surface of the glass slides to be more hydrophobic using a chemical treatment. For this, we pretreated the glass slides with a polydimethylsiloxane (PDMS) solution before the analysis of twitching motility. PDMS is known to coat glass surfaces to make them more hydrophobic (81). As shown in Figure 4B, the treatment of the glass surface with PDMS significantly reduced *P. aeruginosa* twitching to a level that is not significantly different from that on a polystyrene slide. In comparison, PDMS treatment did not impact twitching motility of *P. aeruginosa* on polystyrene slides (Figure 4B), ruling out any inhibitory effects by PDMS. The *P. aeruginosa pilA* mutant showed no twitching under all experimental conditions as expected (Figure 4). These results are consistent with the idea that hydrophilicity of surfaces enhance *P. aeruginosa* twitching, and that bile salts and other detergents stimulate twitching motility on hydrophobic polystyrene surfaces by making them more hydrophilic.

### Increase in hydrophilicity of polystyrene surfaces drastically enhances bacterial twitching motility

While natural polystyrene surfaces are hydrophobic, they can be treated with plasma gas to increase their hydrophilicity for tissue culture (TC) purposes (82). The surfaces of plasma-or TC-treated plates are therefore more hydrophilic than non-treated ones. We compared *P. aeruginosa* twitching motility with 6-well polystyrene plates either TC-treated or non-treated (Figure 5A). The *P. aeruginosa* PAO1 strain exhibited significantly increased twitching motility on plasma-treated surfaces over the non-treated ones in LB media (Figure 5A). The magnitude of increase in this case is about 2-to 3-fold. This increase is more pronounced than on glass surfaces which led to an increase of 1-fold or less (Figures 4A). While the addition of bile salts significantly enhanced *P. aeruginosa* twitching on untreated plates, it had no stimulatory effect on twitching with the TC-treated surfaces.

**Figure 5.**
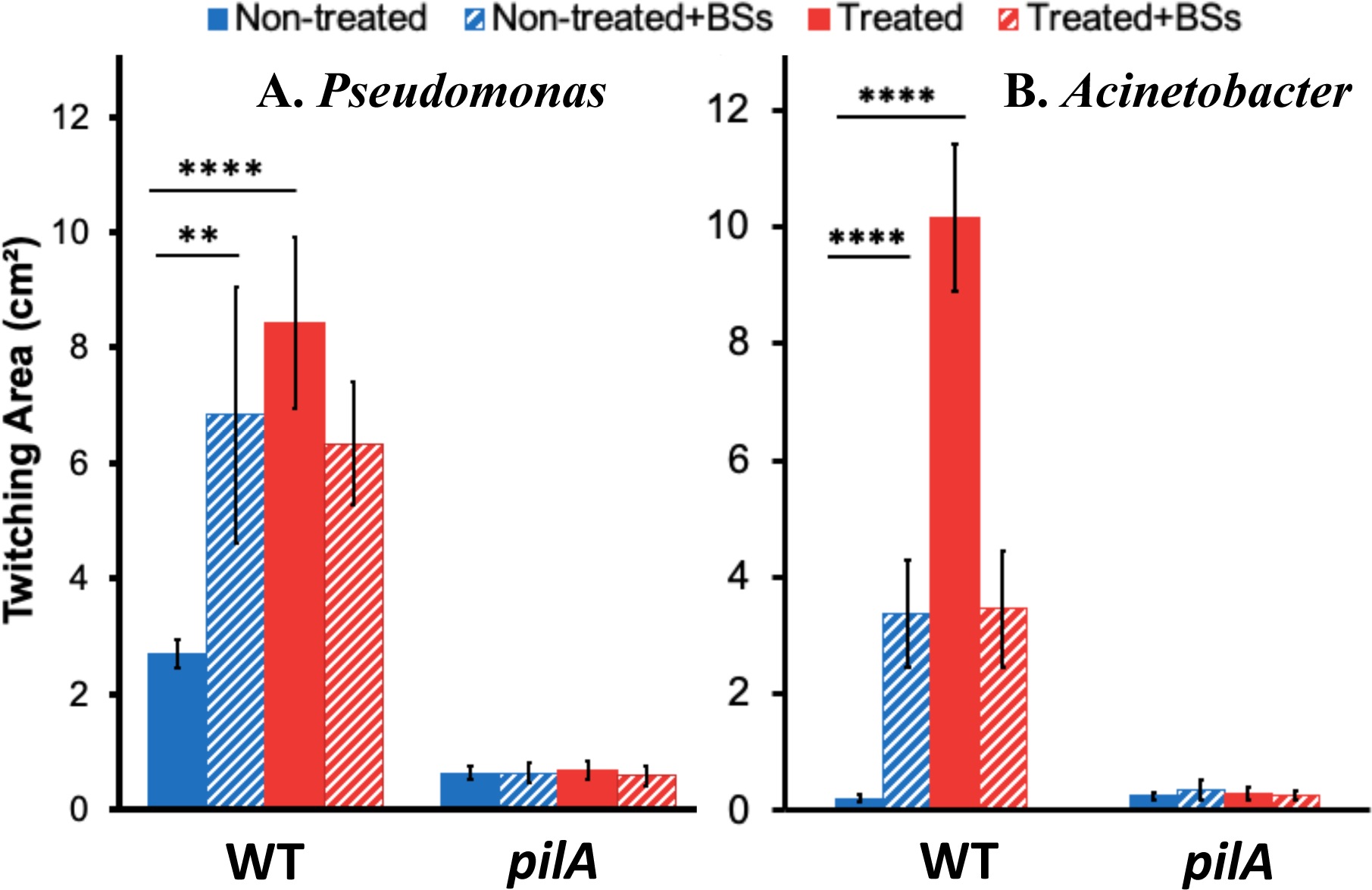
TC-treated plates ameliorate twitching and abolish the effects of bile salts. **A.** *P. aeruginosa* twitching on TC-treated polystyrene. The twitching motility of *P. aeruginosa* PAO1 and its isogenic *pilA* mutant strain was analyzed with 6-well polystyrene plates (see Materials and Methods) either TC-treated or non-treated in LB agar without or with 0.5% bile salts (BSs). **B.** *A. nosocomialis* twitching on TC-treated polystyrene. The twitching motility of *A. nosocomialis* M2 and its isogenic *pilA* deletion mutant was analyzed as in A. Data presented in both panels are from three biological experiments each performed in triplicate. Double (**) and quadruple (****) asterisks indicate that two values are statistically different with P<0.01 and P<0.0001 by the Student’s T-test, respectively.

We examined whether the drastic increase in twitching motility with TC-treated polystyrene surfaces with *P. aeruginosa* could be extended to *A. nosocomialis*. As shown in Figure 5B, *A. nosocomialis* M2 displayed no twitching motility in LB media with the non-treated plates. This is expected because these plates are made of polystyrene like the Petri dishes routinely used for twitching motility assays. TC-treated plates drastically increased M2 twitching with LB media by almost 50-fold without the addition of bile salts or detergents. An isogenic *pilA* mutant was used as a control, and it displayed no twitching under all experimental treatments (Figure 5B). The results here indicate the enhancement of twitching motility by hydrophilic surfaces is not confined to *P. aeruginosa.* Similar enhancement in *A. nosocomialis* suggests a more general phenomenon where hydrophilic surfaces promote interactions that are more favorable for bacterial twitching as mediated by the T4P as a motility apparatus.

## Discussion

The interaction with surfaces is essential for the survival and proliferation of bacteria in their natural environment as well as in health and disease. In their natural habitats, most bacteria exist in multicellular ensembles known as biofilms, the establishment of which depends on bacterial attachment to surfaces (83–86). During bacterial infection of a host, one of the earliest steps in the process is the adhesion of a pathogen to the surfaces of host cells, tissues, and medical implants. From a bacterial perspective, such interactions rely on the timely biogenesis and proper functioning of adhesins on their surfaces. One of the structures critical for bacterial adhesion to both biotic and abiotic surfaces is the bacterial T4P which is prevalent in both Gram-positive and Gram-negative bacteria (2, 13, 28, 29, 31). It is an important virulence factor in many pathogens including *P. aeruginosa* and *A. baumannii* which are both on the WHO priority pathogens list (16). In these bacteria, as well as *A. nosocomialis* and others, the T4P is known to power bacterial twitching motility, which provides a convenient assay for the investigation of T4P biogenesis and function.

Here we described an unexpected mechanism by which bile salts and other detergents can stimulate bacterial twitching motility. It was previously observed that *A. nosocomialis* exhibits twitching motility with MacConkey but not Luria-Bertani (LB) agar media (16). This phenomenon was observed using stab assays to visualize interstitial colony expansion with polystyrene Petri dishes. After confirming this observation, we identified bile salts as the component in MacConkey responsible for eliciting *A. nosocomialis* twitching motility (Figure 1A). The stimulatory effects of bile salts on twitching are not limited to *A. nosocomialis* as we made similar observations with multiple strains of *A. baumannii* (Figure 1B) and *P. aeruginosa* (Figures 2A and S1). Additionally, our results indicated that other detergents, whether anionic or non-ionic, likewise promoted *P. aeruginosa* twitching motility (Figure 3A). Surprisingly, this stimulation of twitching is likely not due to a physiological change in response to the presence of bile salts and other detergents in the growth medium. Instead, it is the ability of detergents to alter the physicochemical properties of a surface that enhances twitching motility in multiple bacterial species.

The above conclusion is based on a few lines of experimental evidence from this study. First, antibiotics with various modes of actions failed to enhance *P. aeruginosa* twitching motility (Figure 3B). These included polymyxin B, which mimics the effects of bile salts and other detergents in applying envelope stress. These results suggested that the enhancement of twitching motility by detergents is likely not a bacterial response to a general or envelope stressor in a growth media. Second, we observed that surfaces of glass, which are more hydrophilic than that of polystyrene, significantly enhanced twitching motility (Figures S2 and 4A). We further demonstrated that the use of glass surfaces abrogated the stimulatory effect of bile salts such that the addition of bile salts no longer promoted *P. aeruginosa* twitching motility on these surfaces (Figure 4A). When the surface properties of glass were changed by the application of a hydrophobic coating, the enhancement of bacterial twitching by glass was reversed (Figure 4B). These results show that hydrophilic surfaces promote twitching, whereas hydrophobic ones suppress it. Because bile salts in the growth media only enhance twitching motility on polystyrene but not on glass surfaces, we conclude that bile salts likely function to modify natural polystyrene surfaces to be more hydrophilic, which promote bacterial motility. Lastly, we performed experiments with TC-treated and nontreated polystyrene surfaces (Figure 5). The TC-treated surfaces, which are more hydrophilic, significantly enhanced bacterial twitching motility. As similarly observed on glass surfaces, the addition of bile salts no longer displayed a stimulatory effect on *P. aeruginosa* and *A. nosocomialis* twitching on TC-treated plates. These results support our conclusion that the physicochemical properties of a surface significantly impact the effectiveness of T4P-powered twitching motility in bacteria. On hydrophilic surfaces, bacteria twitch more, and on hydrophobic surfaces, they twitch less. We further conclude that the promotional effects of bile salts and other detergents on bacterial twitching are largely due to their ability to change the properties of a surface over which bacteria translocate by twitching motility.

Categorically, there are two possible mechanistic explanations for the observed effects of the physicochemical properties of a surface on bacterial twitching. There have been reports that bacteria attach better to hydrophobic surfaces in the context of biofilm formation or otherwise (87–90). It follows that reducing hydrophobicity may lead to alteration of the interactions of a bacterial cell or its pilus with a subsurface over which a bacterium translocates by twitching motility. Sustained twitching movement relies on the recurrence of a multi-step process. These steps include the unobstructed extension of a T4P, the subsequent attachment of the pilus through its distal end for anchoring, followed by a successful T4P retraction event. It is conceivable that tampering with any of these steps through physicochemical interactions with a surface can lead to changes in bacterial twitching behaviors. Alternatively, surface sensing has been demonstrated to mediate changes in cell physiology and behavior (6, 91–96). It is possible that the physicochemical properties of a surface may be detected by a bacterium through surface sensing to modulate T4P biogenesis and its functions, leading to alterations in twitching motility. Further investigation is necessary to determine if the above scenarios or others, either alone or in combination, are the underlying reasons for the observed enhancement of T4P-powered bacterial twitching by surface hydrophilicity.

## Materials and Methods

### Strains and culture conditions

The bacterial strains used in this study are listed in Table 1. These include *A. nosocomialis* M2, *A. baumannii* strains Ab0057 and AYE, and *P. aeruginosa* strains PAO1 and PA14. When appropriate, isogenic *pilA* mutants were used as controls. *P. aeruginosa* strains were grown at 37℃ on 1.5% Luria-Bertani agar (LBA) while *A. nosocomialis* and *A. baumannii* strains were grown at 37℃ on 1.5% MacConkey agar (Oxoid). Oxoid bile acids were used in this study when indicated.

**Table 1.** Bacterial strains used in this study.

| Strain |  | Description | Reference |
| --- | --- | --- | --- |
| *Acinetobacter nosocomialis* | M2 | Clinical isolate | (100) |
| | M2 $\Delta pilA$ | *pilA* deletion and insertion of kanamycin resistant cassette ($\Delta pilA::kan$) | (23) |
| *Acinetobacter baumannii* | Ab0057 | Clinical isolate | (61) |
|  | AYE | Clinical isolate | (62) |
| *Pseudomonas aeruginosa* | PAO1 | Wildtype | (101) |
|  | PW8622 | PAO1 *pilA*-H02::*ISphoA/hah* | (102) |

### Twitching motility assays

Twitching motility was analyzed by three different protocols using the agar stab methods (27) (97) with 1.2% agar media. For all stab assays, the inoculum was prepared by spreading a loopful of bacteria from the outside edge of an overnight plate culture onto a fresh 1.5% LBA plate to create a thin layer of cells. Using a toothpick, bacteria were picked up to inoculate a plate by stabbing through the agar to touch either the polystyrene or glass surface below.

The first protocol uses a standard 100 mm × 15 mm polystyrene or glass Petri dish (Fisher Scientific) as previously described (27, 97). In brief, plates with 25 ml agar media were prepared a day before the assays. These plates were then dried in a biosafety cabinet for 20 minutes before stab inoculation. After 48 hours of incubation at 37℃ in a humidity chamber, the agar media was removed, and the twitching zone was visualized by staining with 1% crystal violet. Twitching areas were determined using the NIH ImageJ software (98).

The second protocol uses either a glass (Opto-Edu) or a polystyrene (VWR International) microscope slide (1″ ξ 3″) inside a standard polystyrene Petri dish for analyzing twitching motility. The microscope slides were first submerged in a filter-sterilized polydimethylsiloxane (PDMS) solution (RainX) (99) or in 70% ethanol as a control. These slides were air dried on a rack at 40℃ before being placed at the bottom of a polystyrene Petri dish. 25 ml of molten agar media was poured over the microscope slides into the Petri dish a day before. A twitching assay is initiated by stab-inoculation as described above except that the incubation is shortened to 18 hours to limit the twitching zone within the boundaries of the microscope slide. One additional modification is that the twitching zone in this case was visualized by incident light and traced with a permanent marker without removing the agar from the Petri dish. Twitching area was determined using ImageJ as above.

The third protocol uses 6-well polystyrene plates with or without TC-treatment (Falcon). Each well of a plate contains 2 ml agar media and twitching motility assays were initiated by stab inoculation as described above. The twitching area was analyzed as described for the microscope slide-based assay except that the incubation time was extended to 24 hours.

## Acknowledgements

The research in the Yang laboratory is partially supported by the National Science Foundation grants MCB-1919455 and EFMA-2318093, as well as the CHRB award 208-07-23 to Z.Y. The Yang laboratory additionally received a pilot grant from the Center for Emerging, Zoonotic and Arthropod-borne Pathogens. M.T.O. was the recipient of the Presidential Scholarship Initiative from Virginia Tech and the David Lyerly Undergraduate Microbiology Research Fellowship from the Department of Biological Sciences at Virginia Tech. K.J.D was the recipient of a GSDA and the Lewis Edward Goyette Fellowships as well as the Liberati Scholarship from Virginia Tech.

M.T.O., K.J.D., and Z.Y. designed research and analyzed data. M.T.O. and Z.Y wrote the manuscript. M.T.O., T.M.S., and D.H. performed experiments. T.M.S. aided in figure preparation and protocol development.

We thank Dr. Kurt Piepenbrink, Dr. Robert Bonomo, Dr. Helen Zgurskara, and Dr. George O’Toole for providing bacterial strains. We additionally thank Ghazal Soleymani for proof-reading the manuscript.

There is no conflict of interest to declare.

