## Supplemental Materials for "Surface Hydrophilicity Promotes Bacterial Twitching Motility"

**Table S1. Ingredients of LB and MacConkey agar media for twitching.**

| **MacConkey (MC) agar*** | **Luria-Bertani (LB) agar*** |
|---|---|
| 1.2% Agar | 1.2% Agar |
| 0.5% Sodium Chloride | 0.5% Sodium Chloride |
| 0.075% Neutral Red | 0.5% Yeast Extract |
| 2% Peptone# | 1% Tryptone |
| 1% Lactose# | |
| 0.5% Bile Salts# | |

*Ingredient concentrations are in % (w/v).

### Ingredients highlighted in blue signify key differences in media composition.

**Table S2. Properties of detergents**.

| Detergent | MaxNIC (μg/mL)* | CMC (mM)** | Type** |
|---|---|---|---|
| **SDS** | 750 | 7-10 | Anionic |
| **Triton-X100** | 75 | 0.23 | Non-ionic |
| **Triton-X114** | 75 | 0.2 | Non-ionic |

* The maximum non-inhibitory concentration (MaxNIC) of each detergent against *P. aeruginosa* PAO1 was determined experimentally in liquid growth media in the presence of detergent at different concentrations (data not shown).

** CMC (critical micelle concentration) and types of detergent were from the reference data from Sigma-Aldrich (Bhairi et al., 2017).

**Table S3. Antimicrobial susceptibility testing of *P. aeruginosa* strain PAO1.**

| Antibiotic | MaxNIC (ng/ml)* | Target |
|---|---|---|
| Ampicillin | 313 | Cell wall synthesis |
| Ciprofloxacin | 31 | DNA gyrase |
| Gentamicin | 31 | 30S ribosomal rRNA |
| Polymyxin B | 313 | Outer membrane |

* The maximum non-inhibitory concentration (MaxNIC) of each antibiotic against *P. aeruginosa* PAO1 was determined experimentally by liquid growth media in the presence of antibiotic at different concentrations (data not shown).

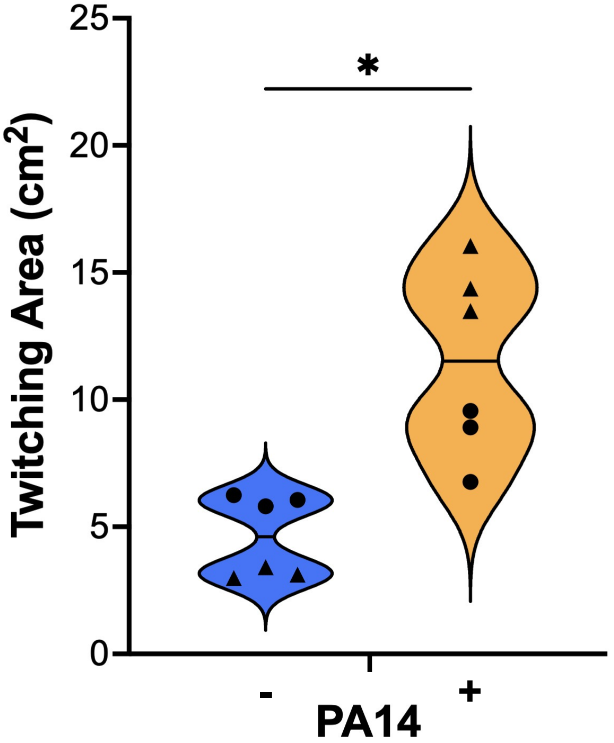


**Figure S1. Twitching motility of *Pseudomonas aeruginosa* PA14 in the presence of bile salts.** Twitching motility was analyzed and presented as in Figure 1B. Data presented is from two biological experiments each performed in triplicate. An asterisk (*) specifies that the twitching area values with bile salts for the indicated strains are statistically different ($P<0.05$) from those without bile salts.

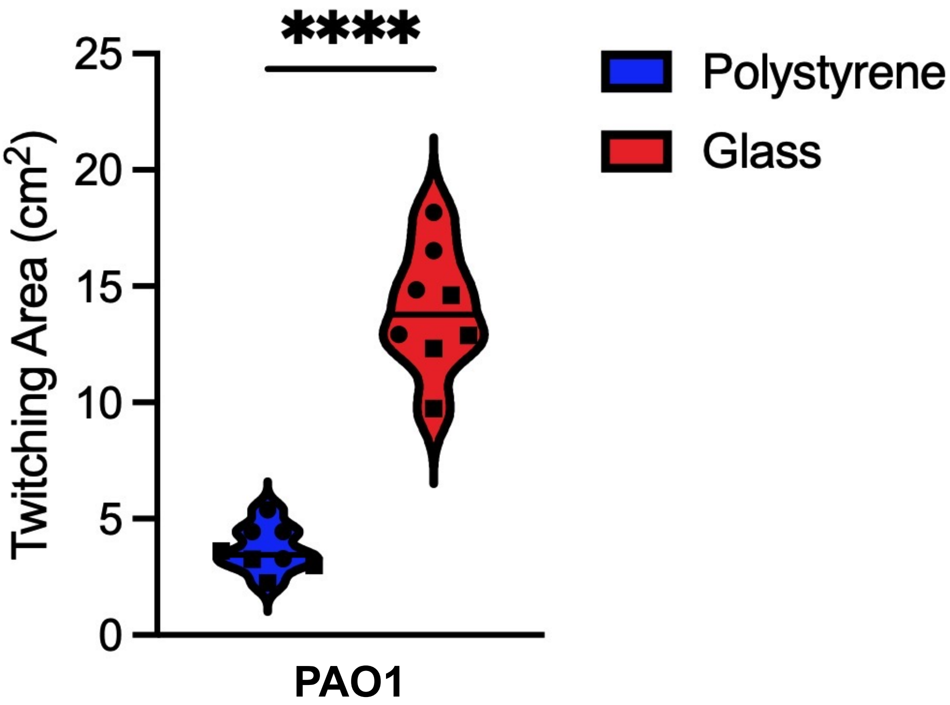


**Figure S2. Glass surfaces enhance the twitching motility of *Pseudomonas aeruginosa*.** Twitching motility of *P. aeruginosa* PAO1 were analyzed on polystyrene or borosilicate glass Petri dishes in Luria-Bertani (LB) agar, as in Figure 1B. Data presented is from two biological experiments each performed in quadruplicate.
